# Actomyosin, vimentin and LINC complex pull on osteosarcoma nuclei to deform on micropillar topography

**DOI:** 10.1101/822445

**Authors:** Nayana Tusamda Wakhloo, Sebastian Anders, Florent Badique, Melanie Eichhorn, Isabelle Brigaud, Tatiana Petithory, Maxime Vassaux, Jean-Louis Milan, Jean-Noël Freund, Jürgen Rühe, Patricia M. Davidson, Laurent Pieuchot, Karine Anselme

## Abstract

Cell deformation occurs in many critical biological processes, including cell extravasation during immune response and cancer metastasis. These cells deform the nucleus, its largest and stiffest organelle, while passing through narrow constrictions *in vivo* and the underlying mechanisms still remain elusive. It is unclear which biochemical actors are responsible and whether the nucleus is pushed or pulled (or both) during deformation. Herein we use an easily-tunable poly-L-lactic acid micropillar topography, mimicking *in vivo* constrictions to determine the mechanisms responsible for nucleus deformation. Using biochemical tools, we determine that actomyosin contractility, vimentin and nucleo-cytoskeletal connections play essential roles in nuclear deformation, but not A-type lamins. We chemically tune the adhesiveness of the micropillars to show that pulling forces are predominantly responsible for the deformation of the nucleus. We confirm these results using an in silico cell model and propose a comprehensive mechanism for cellular and nuclear deformation during confinement. These results indicate that microstructured biomaterials are extremely versatile tools to understand how forces are exerted in biological systems and can be useful to dissect and mimic complex in vivo behaviour.

## INTRODUCTION

Cells reside in intricate 3 Dimensional (3D) microenvironments within which they are subjected to various mechanical constraints that modulate their behavior. The nucleus, which is the largest and stiffest organelle within the cell, represents a physical constraint that limits cell migration in narrow spaces.(1-4) The impact of cell confinement on nuclear deformation and cell mechanics have been extensively studied using physical methods such as micropipette aspiration, micro-indentation or isolating nucleus from cells.(5-8) Although these techniques have established some new paradigms on cell-nuclear mechanics, they do not reflect how cells spontaneously deform *in vivo*. Microfabrication methods have emerged as an extremely versatile tool to engineer and recapitulate various microenvironments, allowing direct visualization of cell spontaneous confinement.(9-12) Microfluidic channels have proved very useful to study cell migration. Using these, actin, vimentin and linker of the nucleoskeleton and cytoskeleton (LINC) complex proteins have been implicated in translocating the nucleus through constrictions during migration.(3) Microtubules have also been implicated in nucleus movement in neurons and muscle cells.(13, 14).These studies indicate that the nucleus may be pushed from behind or pulled from the front, although it has remained difficult to precisely define the mechanism.(15)

We previously demonstrated nuclear deformation *ex situ* on 3D micron-sized pillars (16-18): osteosarcoma cells spontaneously remodel their nuclei to fit within the available spaces between 3D micropillars whereas healthy bone cells span the tops of the pillars. (19) These pillar arrays were further used by other researchers to evaluate various cell lines for their ability to deform, or the impact of confinement on differentiation processes.(20-23)

We show here that our micropillar arrays offer an elegant and simple method to dissect the deformation behaviour of cells in 3-D constrictions. This assay allow us to study nuclear deformation caused by the cell, independently from cell migration, in an assay in which the surfaces can be chemically modified and the cells can easily be treated with drugs and collected for biochemical studies. We combine 3D micropillar topography of controlled surface chemistry with live imaging and *in silico* cell numerical simulation to understand the mechanism by which cells deform their nuclei. We show that nuclear deformation of osteosarcoma nuclei in confinement is orchestrated by pulling forces which are regulated by actomyosin coupled to LINC complex proteins and is assisted by vimentin intermediate filaments.

## MATERIALS AND METHODS

All products were purchased from Sigma–Aldrich (France) unless specified.

## CELL CULTURE

SaOs-2 osteosarcoma cells (ATCC, USA) were cultured in a humidified incubator at 37 °C and 5% CO_2_. McCoy’s 5A Modified medium was supplemented with streptomycin (0.1 mg/ml), penicillin (100 U/ml), heat inactivated fetal bovine serum (15% v/v, Biowest, France) and L-Glutamine (2mM). For all experiments 1X10^5^/ml cell density was used in 60mm petri dishes.

## MICROPILLAR FABRICATION, CELL SEEDING ON PILLAR TOPOGRAPHY, CHEMISTRY MODIFICATION

For more details, see previous publications (16, 17, 19)

### PDMS CAST

silicon templates with square micropillars measuring 7µm (height) x 7µm (length) x 7µm (space) were fabricated in IMTEK (Freiburg, Germany). Polydimethylsiloxane (PDMS) (Sylgard 184, Dow Corning, USA) negative replica stamps were produced by uniformly mixing the precursor and curing agent in 10:1 ratio and pouring it over the silicon template glued in an aluminum cup. Following degassing in the vacuum pump for 30 minutes to remove air bubbles, PDMS mixture was cured at 60°C in an oven overnight (ON). Hardened PDMS was then separated from template carefully and used to prepare Poly-L-Lactic acid (PLLA) micropillars.

### PLLA FILM

PLLA powder (Evonik, Germany) was mixed in chloroform solution and poured in glass petri plate. Uniform PLLA concentration and volume were chosen to keep it constant per unit area of the 10cm glass Petri plate (0.002g/cm^2^). PLLA film was obtained by allowing the mixture to air dry in a fume hood overnight and then cut into pieces for further use.

### PILLAR FABRICATION

Hot embossing was prepared by heating the PDMS negative replica above the glass transition temperature of PLLA (180°C). The piece of PLLA film was placed over the PDMS template and pressed down manually for approximately 5-10 sec. The assembly containing PDMS-PLLA sandwich was quenched in cold water to vitrify PLLA before demolding.

### SAMPLE PREPARATION

Samples were glued using cell compatible aqua-Silicone glue (Den Bravent Sealants, Netherlands) in 60mm Petri plates and then sterilised using 70% ethanol for 10 minutes and dried under laminar air flow.

### MICROPILLAR DIMENSIONS AND CHEMICAL MODIFICATIONS

Polymeric micropillars were fabricated by double replication. As an original, a structured Si wafer was used. The wafer was microstructured with square micropillars with a width of 10 µm, a height of 10 µm, and spacing of 3 µm. The Silicon wafer was made by the clean room service center of the University of Freiburg (IMTEK). For the first replication step (negative), a commercial PDMS elastomer kit (Sylgard 184, Dow Corning, Midland, USA) was used; base and curing agent were mixed in the ratio of 10:1 (w/w). The mixture was poured over the etched Si wafer and cured at 80 °C for 3 h. The hard PDMS was peeled off the Silicon wafer. The polymer used in the second replication step (positive copy) was obtained by free radical polymerization of n-butyl acrylate (nBA) and 4-(methacryloyloxy) benzophenone (MABP); 5 mol% MABP were used. The resulting P(nBA-co-5%MABP) was dissolved in toluene (100 g/L). A drop of the dissolved polymer was placed on a glass slide (Menzel Gläser cover glasses, round 18 mm, No. 3, VWR, Germany). The PDMS stamp was pressed on top of the dissolved polymer. The combination of all three components (stamp, polymer and glass slide) was crosslinked by UV irradiation (λ = 365 nm, 12.98 J, 90 min). The PDMS stamp was peeled off to liberate the completed microstructured surface. The surface was modified by different methods: To modify the entire surface, spin coating (1000 rpm, 60 sec) with polymer solutions was used either with the cell-repellent (Poly dimethyl acrylamide) P(DMAA-co-5%MABP) (solution in ethanol, 50 g/L), or the cell-attractive (AA-all attractive) P(nBA-co-5%MABP) (solution in toluene, 50 g/L). To modify only the top of the pillars (TA-top attractive, SA-space adhesive), a plane aluminum foil (1 cm× 4 cm) was dip coated (immersion and retraction speed 100 mm/min) with the dissolved polymer. The aluminum foil was then pressed on top of the microstructure and promptly removed. The coated polymer was attached to the surface by UV light crosslinking (λ = 365 nm, 12.98 J, 90 min). For confocal verification of coatings, cell adhesive polymer was tagged with Cyn-5 streptavidin (GE Healthcare, USA) at a 1:500 dilution and cell repellent polymer was tagged with Rhodamine B isothiocyanate (Sigma, France) at 1% concentration with the polymer.

## DRUG TREATMENT

Concentrations were used as follows: Cytochalasin D (1 µM), Latrunculin B (2 µM), Colchicine (5 µM), Nocodazole (10 µM), 1-(5-Iodonaphthalene-1-sulfonyl)-1H-hexahydro-1,4-diazepine hydrochloride [*ML 7*] (20 µM), Blebbistatin (25 µM), Acrylamide (10 mM) and Iminodipropionitrile [*IDPN*] (2%). The optimum concentration (more than 50% viability) was determined using an MTT colorimetric assay 6 hours after drug treatment and measuring the resulting absorbance at 570nm (normalised at 620nm) using an EZ READ 400 Plate reader (Bichrom, UK). Percentage viability graph was plotted (**S5).** Drugs were added in the medium containing cells in suspension and then seeded on pillars for 6 hours. Drug washout experiments were performed at 2, 5 and 20 hours with1X PBS (Gibco, Life technologies, USA).

## IMMUNOSTAINING

Unless specified products were purchased from Sigma Aldrich (France). Cells were fixed with 4% formaldehyde (Electron Microscopy Sciences, USA) and permeabilized using 0.5% Triton X-100 for 15 min and blocked with 3% bovine serum albumin for 1 hour. Incubation with primary antibodies was done for 1 hour with following antibody concentration: anti-vimentin (1:50; V6630; anti-Mouse), anti-βtubulin (1:200; T4026; anti-Mouse), anti-myosin IIA (1:100; M8064; anti-Rabbit), anti-paxillin (1:250; ab32084; anti-Rabbit; Abcam-UK), anti-pericentrin (1:100; ab4448; anti-Rabbit; Abcam-UK), anti-Histone deacetylase 1 (1:200; H3284; anti-Rabbit), anti-Sun 1 (1:100; M8064; anti-Rabbit), anti-Sun 2 (1:100; M8064; anti-Rabbit), anti-SYNE2 (1:50; HPA003435; Atlas, Sweden), anti-Lamin A, specific for lamin A only and not lamin C (1:500; L1293; anti-Rabbit). After short rinses in 1X PBS, secondary antibodies were incubated for 1 hour using Alexa 488 (1:200; ab150073; anti-Rabbit; Abcam-UK), Alexa 488 (1:200; ab150113; anti-Mouse; Abcam-UK), Alexa 647 (1:200; ab150115; anti-Mouse; Abcam-UK). Confocal acquisitions were obtained after PBS rinses. Actin-555 (1:20; A34055; Thermoscientific-France) or phalloidin-FITC (Fluorescein-5-isothiocyanate) (0.4µg/ml) were used to label actin and Hoechst (1:1000; 33258; Thermoscientific-France) was used to label nucleus.

## TRANSFECTION

Dominant Negative-KASH plasmid DNA was kindly gifted by Dr Nicolas Borghi (Jacques Monod Institute, Paris-France). Lamin A-GFP was used to overexpress lamin A (17653, Addgene, USA). Cells cultured up to 60-70% confluency were transfected with plasmids using Lipofectamine 3000 (Thermoscientific, France) upon manufacturer’s recommendations. Culture media used for transfection was deprived of antibiotics. After transfection, cells were seeded for 48-72 hours on pillars before imaging.

For the nuclear deformation of the cell over time, cells were transfected with baculovirus kit for actin-RFP and histone-GFP (BacMan 1.0 and 2.0, Thermoscientific-France). The transfection was carried out according to the manufacturer’s protocol. The particle per cell (PPC) value was determined to be 40 for SaOS-2. For histone-GFP kit 1.0, cells were incubated for 4 hours with baculovirus, followed by 2 hours incubation with enhancer at 37 °C with 5% CO_2_ in a humidified incubator. For actin-RFP kit 2.0, the cells were directly transfected with the baculovirus and allowed to grow for 16 hours at 37 °C with 5% CO_2_ in a humidified incubator.

## SiRNA TREATMENTS AND QUANTITATIVE REAL TIME-PCR

SiRNA (Thermo Scientific): non-target negative control (AM4613), *SYNE2* (s23330), *SUN2* (s24465), SUN1 (s23629) and *LMNA* (s8221). Transfection was carried out using Lipofectamine® RNAiMAX Transfection Reagent (Thermoscientific, France) following the manufacturer’s instructions. 10pmol concentration of siRNA was selected for the knockdown. Vimentin siRNA was used at a 40pmol concentration (sc-29522, Santa Cruz Biotechnology, USA). RNA extraction was carried out after 72 hours of KD treatments using RNAeasy micro-kit (Qiagen, France). The quantity and quality of RNA was determined using a Nanodrop spectrophotometer (Maestro, Taiwan). Retrotranscription was performed using iQ™ SYBRgreen mastermix (Biorad, Switzerland) in iScript cDNA synthesis instrument (Biorad, Switzerland). The data was normalized against the 18S methyltransferase *(emg1)* housekeeping gene. Primers were designed with melting temperature at 60°C, using the GenScript Real-time PCR Primer Design algorithm (https://www.genscript.com/tools/real-time-pcr-tagman-primer-design-tool). The qRT-PCR efficiency was determined using LinRegPCR software (Heart failure research centre, Netherland). Expression levels were further analyzed based on ΔΔCT method (n=2). The primer sequences are given in **S8.** The qPCR program was one cycle at {95 C-30 s}, followed by 39 cycles at {95 C for 5 s} and {60 C for 30 s}. PCR products were then sequentially heated to {65 C for 2 s} and to {95 C for 5 s} to measure the dissociation curve. Experiments were independently repeated at least twice. qPCR efficiency was assessed using the LinRegPCR program (version 2015.2). The expression levels of the genes were analyzed based on the ΔΔCT method, as described by Livak et al.(39)

## WESTERN BLOT ANALYSIS

Unless specified, instruments and supplies were all purchased at Bio-Rad (Germany). Cell lysis and protein resuspension were achieved in radio immunoprecipitation assay (RIPA) buffer. Total protein amount was estimated using the colorimetric Micro BCA™ Protein Assay (Thermoscientific-France). 20μg of total protein sample was loaded on a precast Mini-protean TGX stain-free gel and separated according to size by electrophoresis. Protein gel images for normalization were acquired using ChemiDoc™ Imaging Systems before transfer. Proteins were transferred from the gel to 0.2μm polyvinylidene difluoride (PVDF) pre-activated membranes using the Transfer Blot turbo system. Afterwards, unspecific sites were saturated with 3% BSA (Sigma, France) for 1 hour at room temperature (RT). Primary antibody incubation was performed overnight at 4°C, for 1h at RT. Primary antibody incubation was performed overnight at 4°C, using the following antibody concentrations: lamin A (1:500, L1293, Sigma, France), vimentin (1:200; V6630; anti-Mouse; Sigma-France), SUN1 (1:1000; ab124770; anti-rabbit; Abcam-UK), SUN2 (1:100; lot A45573; anti-rabbit; Sigma-France) and nesprin2 (1:1000; ab124770; anti-rabbit; Abcam-UK). After short rinses, blots were incubated with the Horse Radish Peroxidase (HRP) Goat anti-rabbit IgG-HRP conjugate (1:3000, Cat No. 170-6515) and Goat anti-mouse IgG-HRP conjugate (1:3000, Cat No. 170-6515) for 1h at RT. Protein bands were revealed using a chemiluminescent substrate (Clarity™ Western ECL Blotting Substrates) and digitally acquired using the ChemiDoc™ Imaging Systems. Full-sized immunoblots images are given in **S9.**

## CONFOCAL IMAGING

Imaging was performed using an upright Carl Zeiss LSM 700 confocal microscope, Germany, using ZEN software. For live imaging z stack were acquired using 63X/1.4 oil; 63X/1.0 VIS-IR water or 20X/1.0 DIC VIS-IR W Plan-Apochromat objective (Zeiss, Germany) equipped with a temperature (Okolab, Italy), CO_2_/O_2_ (Okolab, Italy), and humidity controlled chamber.

## IMAGE ANALYSIS AND QUANTIFICATION

### NUCLEAR DEFORMATION

3-D images of nuclei were quantified using ImageJ/Fiji (NIH, version 2.0.0-RC-61/1.51n, USA).(40) All images to quantify nuclear deformation, were taken via live cell imaging in order to minimize artefacts due to fixation. The area below the pillars was considered as deformed and above pillars was considered undeformed. The deformed area was divided by the total cross-sectional area of the nucleus to obtain the percentage of deformation (see **S1).**

### FOCAL ADHESION LOCALIZATION ON PILLARS

The height, the number of slices and the interval between slices was kept constant for all independent acquisitions. The image was then thresholded and the resulting particles were analyzed. The average number of FA in each section was plotted on the Y-axis showing the height of pillars and X-axis showing the average number of FA/cell.

## *IN SILICO* CELL MODEL

The cell model was developed and validated previously to analyze the influence of the substrate convexity on cell adhesion and nucleus strain in a previous study.(41) The in silico cell model is extensively described in Vassaux et al.(30) In the present study, the only modification to the model was to add the substrate composed of micropillars. The composition of the model and the adhesion process are same as used in Vassaux et al.(41)

## STATISTICAL ANALYSIS

All the results were checked for significance by using one way ANOVA test followed by two-tailed Student’s *t*-test (unequal variance) for pairwise comparison and Dunnet’s test for multiple comparisons, using GraphPad Prism software (GraphPad Software, Inc). P-values were calculated by two-tailed unpaired Student’s t-test as compared to control untreated cells for comparison between two groups and Dunnett’s multiple comparisons test for comparison between more than two groups. The significance value obtained were categorized as P > 0.05 for ns (not significant). The error bars represent the standard deviation of the measurements. All experiments were performed independently three times.

## RESULTS

### Nuclear deformation is concomitant with chromatin, cytoskeleton and focal adhesion reorganization

#### Kinetics of nuclear deformation and chromatin reorganization

We previously showed that SaOs-2 cells deform extensively on micropillars. Here we focus on this cell line and investigate the kinetics of nuclear deformation **(S1 and method section)** and cell adhesion on micropillars. Deformation of the nucleus is detectable one hour after the cell contacts the surface. The nucleus is progressively inserted in the interpillar space, reaching a maximal and steady deformation at 24 hrs (**Fig. 1a,b** and **movie SM1**). Here we define nuclear deformation percentage as the proportion of the nucleus below the tops of the pillars. (See Methods for more details.) Unless otherwise mentioned, all nuclear deformation measurements were performed after 24 hrs of incubation.

**Figure 1.**
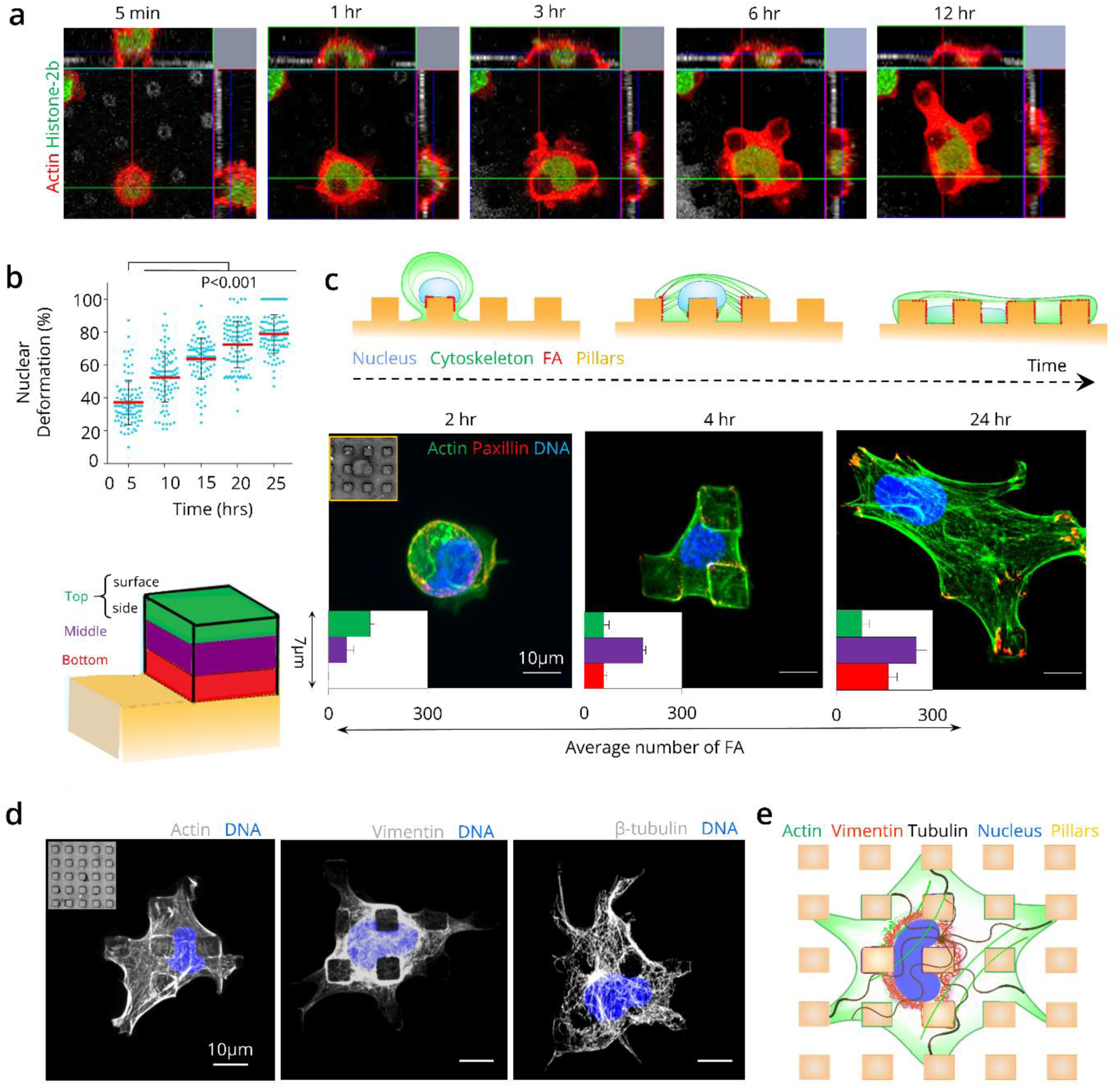
Nuclear deformation involves focal adhesions, chromatin and cytoskeletal dynamics. **a)** SaOs-2 deformation kinetics on pillars. Orthogonal images show cells expressing actin-RFP (red) and histone H2B-GFP (green) seeded on micropillars at different time points during cell adhesion (for video see supplementary movie 1). **b)** Quantification of nuclear deformation. (For a description of the analysis see supplementary figure S1). The mean value is indicated by a red line. P-values compared to initial cell adhesion on pillars at 2 hours. (n=150/time point, 3 independent experiments.) **c)** FA localization and nuclear deformation on pillars, z-stack projection of SaOs-2 cells (actin-green; paxillin-red; nucleus DNA-blue) at different time points. Subsets represent the average number of FAs at different heights (see figure on left). (n=45/time point, 3 independent experiments). **d)** Immunostaining (vimentin & β-tubulin) and phalloidin (actin) labelling showing cytoskeleton elements in grey. **e)** Schematic summary of the cytoskeletal organization after nuclear deformation on pillars. All error bars represent ±SD.

We observed that cells increased in number over time within the pillar arrays, which raised the question of cell division in confined cells. We found that the cells avoid confinement during division and ascend perpendicularly to the micropillar substrate to perform mitosis **(S2 & SM 2)**. The daughter cells subsequently invade the structure quickly after cytokinesis, faster than the initial deformation, likely because the daughter cells retain focal adhesions from mother cells that aid in spreading (24).

#### Focal adhesion localization during nuclear deformation

Cells adhere to surfaces using integrin mediated focal adhesions (FA). We analyzed the FA localization on pillars at different timepoints **(Fig. 1c)**. To facilitate the quantification, we segmented the pillars into three zones (top, middle and bottom) and quantified the average number of FAs for each area (see methods for details). FAs localize initially at the top part of pillars and gradually spread on the lateral sides. FA density increases over the initial 6-8 hours and reaches a plateau, concomitantly with the nuclear deformation kinetics. Nuclear deformation progression is correlated with an increase in FA density on the mid-lateral regions of the interpillar space.

#### Cytoskeletal organization during nuclear deformation

The cytoskeleton plays a crucial role in adapting the cell to its environment. We thus looked at the organization of cytoskeletal components (microfilaments, intermediate filaments (IFs) and microtubules) in deformed SaOs-2 cells. We observed strong accumulation of actin on the edges of pillars and uniform distribution throughout the cell with an increase in stress fibers across the top of the cell after 24 hours. IFs exhibited higher density around the pillars and accumulated around the nucleus at the center of the cell. Microtubules were uniformly spread throughout the cell **(Fig. 1d).** The cytoskeleton and FA organization during nuclear deformation is summarized in **Fig. 1e**.

### Nuclear deformation is governed by actomyosin and intermediate filaments dynamics

We dissected the contribution of microfilaments (actin), myosin, IFs (vimentin) and microtubules (β-tubulin) using cytoskeletal disrupting drugs. Optimal drug concentrations were determined using a cell viability assay (MTT) to determine the concentration at which at least 50% of cells survived drug treatment **(S3 and methods).** We used the percentage of nuclear deformation as a readout for nuclear confinement. We found that depolymerizing actin (cytochalasin D, latrunculin B), inhibiting the actin motor protein myosin II activity (blebbistatin, ML7) as well as depolymerizing IFs (acrylamide, IDPN) strongly reduced nuclear deformation compared to control cells **(Fig. 2a and b).** Microtubule depolymerization (colchicine, nocodazole) did not have any effect on nuclear deformation. Because acrylamide can also affect actin and microtubule organization (25), we confirmed the results using a gene silencing technique (RNA interference) for vimentin IFs specifically. The silencing efficiency was confirmed by RT-qPCR and western blot techniques **(Fig. 2c and d).** Nuclear deformation after vimentin knockdown (KD) confirmed the role of IFs in assisting nuclear deformation on micropillars **(Fig. 2e).** We performed drug washout control experiments for actin, myosin and vimentin. Our results showed that nuclear deformation was regained over time (2, 5 and 20 hours) after the respective drugs were washed out **(S4).** Our results indicate that acto-myosin and IFs play crucial roles in cell-nuclear confinement, but targeting microtubules does not affect nucleus deformation.

**Figure 2.**
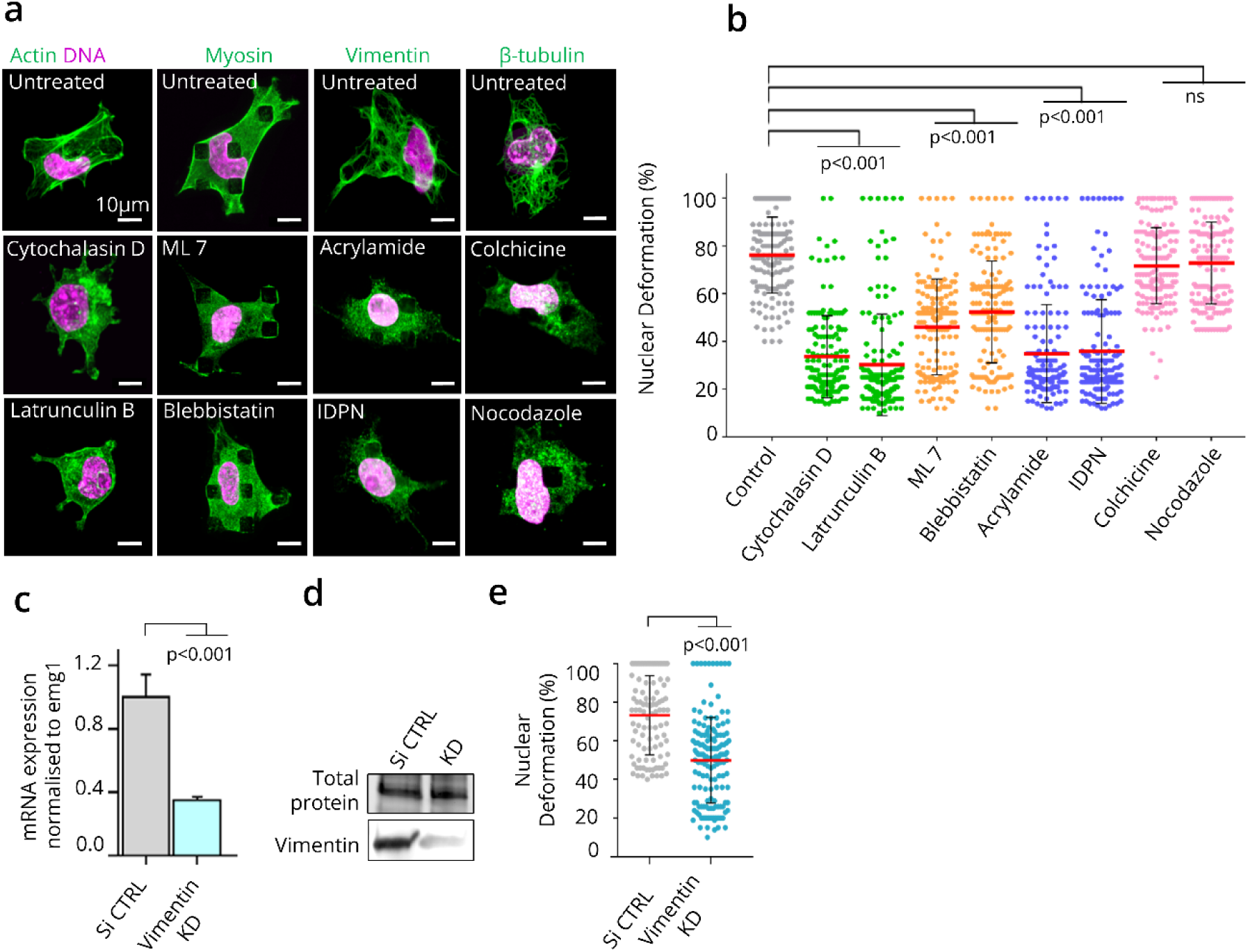
Actomyosin and IF drive nuclear deformation. **a)** Immunostaining of SaOs-2 cells on micropillars 24 hrs after drug treatments. Z stack projection of cells immunostained with actin (green) and nucleus (magenta); myosin IIA (green); vimentin (green), β-tubulin (green). **b)** Quantification of nuclear deformation of SaOs-2 6 hours after drug treatment (n=160/drug treatment) with mean (red line). **c)** RT-qPCR was used to access vimentin expression in knockdown and non-target control (non-target RNA) cells (grey) normalised to emg1 gene expression, after 48 hours. **d)** Western blot was used to access vimentin protein levels after 48 hrs in knockdown and control cells. **e)** Quantification of nuclear deformation in vimentin knockdown and non-target control cells, after 48 hours, (n=150 nuclei/siRNA treatment). All experiments were performed independently three times.

### Coupling the cytoskeleton to the nucleus is necessary for nuclear deformation

#### The LINC complex is required for nuclear deformation

The nucleus is connected to the cytoskeleton by nuclear envelope proteins which form the LINC complex. (26) Cytoskeletal components (actin, microtubules, IFs) are connected to nesprins, which cross the outer nuclear membrane, and in turn are connected to SUN proteins which cross the inner nuclear membranes. We silenced SUN1, SUN2 and SYNE2 (nesprin2) individually and confirmed the silencing efficiency by immunofluorescence, RT-qPCR and western blot **(Fig. 3a-c).** Our results showed a small, but non-significant reduction in nuclear deformation in KD cells compared to control cells **(Fig. 3d).** Functional redundancy within SUN and Nesprin variants that were not silenced might have played a compensatory role in the process. We attempted to target multiple LINC components at the same time using the same RNAi approach but the silencing efficiency was reduced significantly.

**Figure 3.**
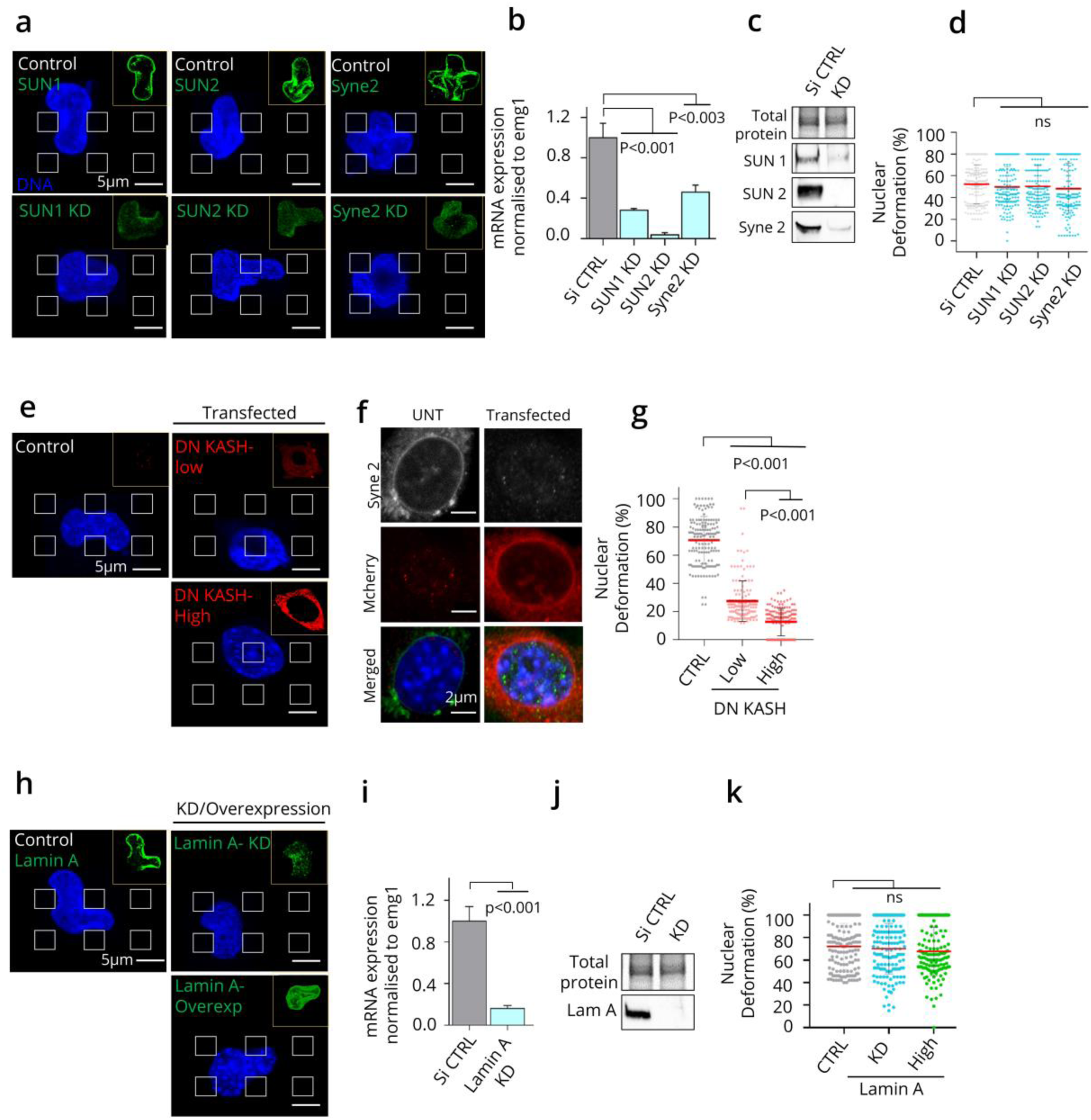
Nuclear deformation requires LINC-cytoskeletal coupling. **a)** SaOs-2 nuclei on micropillars 24 hours after siRNA treatments. Immunostaining for SUN1, SUN2 and Nesprin2 (SYNE2) at right top corner and Hoechst staining for nucleus (blue). All images are Z stack projection and grey squares represents the position of top part of pillars. **b)** RT-qPCR was used to access SUN1, SUN2 and SYNE2 expression in knockdown and control cells (grey) normalized to emg1 gene expression. **c)** Western blot was used to access SUN1, SUN2 & syne2 protein levels in knockdown and control cells. **d)** Quantification of nuclear deformation after 48 hrs of siRNA treatment (n=150/SiRNA treatment) showing mean (red line). **e)** Confocal images of SaOs-2 transfected with DN-KASH mCherry showing low and high DN-KASH expression. **f)** Displacement of endogenous nesprin (grey) after DN-KASH transfection compared to untreated cells. **g)** Quantification of nuclear deformation 48 hours after transfection of DN KASH in cells presenting high, low and no (CTRL) mCherry signal (n=150/SiRNA treatment). The mean value is shown with a red line. **h)** Confocal images of SaOs-2 after 48 hrs of lamin A knockdown (green) and lamin A-GFP overexpression (green). **i)** Lamin A/C expression in knockdown and control cells (grey) evaluated by RT-qPCR. **j)** Lamin A protein levels in knockdown and control cells. **k)** Graph showing nuclear deformation percentage after 48 hrs of lamin A knockdown, high lamin A-GFP overexpression and control cells (grey), (n=150/SiRNA treatment) with mean (red line). ‘n’ is number of cells analyzed over three independent experiments. P-values were calculated by two-tailed unpaired Student’s *t*-test as compared to control untreated cells for comparison between two groups and Dunnett’s multiple comparisons test for comparison between more than two groups. All error bars represent ±SD; ns, not significant.

Nesprins have a highly conserved KASH (Klarsicht, ANC-1, Syne homology) domain at their C-termini that binds SUN proteins while their N-termini associate with different cytoskeletal constituents. Thus, we disconnected the LINC complex by overexpressing a dominant-negative KASH domain (DN-KASH-mCherry) **(Fig. 3e and 3g**), resulting in displacement of endogenous nesprins to the ER.(27) Nesprin 2 immunostaining confirmed the delocalization of endogenous nesprins after DN-KASH overexpression **(Fig. 3f)**. Cells overexpressing DN-KASH deformed their nucleus less than control cells **(Fig. 3g)**. Strikingly, the effect was dose-dependent: cells accumulating the highest DN-KASH levels presented the lowest nucleus deformation **(Fig. 3g)**. Altogether, these results highlight the central role of LINC-cytoskeleton coupling in nuclear deformation and cell confinement.

#### Modulation of Lamin A levels does not affect the ability of SaOs-2 cells to deform their nuclei

Lamin A proteins play a central role in the regulation of nuclear mechanics.(1, 28, 29) We therefore tested whether modulating their levels has an effect on SaOs-2 nuclear deformation. We downregulated endogenous lamin A/C using siRNAs or overexpressed a lamin A-GFP construct **(Fig. 3h-k)**. Surprisingly, neither the increase nor decrease in A-type lamin levels affected the nuclear deformation ability at 24 hours **(Fig. 3k)**. A-type lamins therefore do not determine the ability of cells to deform their nuclei.

### Cell adhesion to the lateral sides of the pillars is necessary for nuclear deformation

We investigated how the cells interact with the substrate to deform themselves to understand whether they attach and pull or push from above to squeeze themselves in narrow spaces. We created three types of surfaces: (1) all adhesive pillars, (2) top adhesive pillars and (3) interspace adhesive pillars, in which the tops of the pillars are cell-repellent but the cells can adhere to the space in between the pillars (**Fig. 4 a**). We used a more restrictive spacing (3µm) to obtain an intermediate level of deformation and detect increases as well as decreases in the extent of deformation. On “all adhesive” pillars, nuclear deformation at 24 hours reached 30±7% (**Fig. 4 b**). In contrast, on “top adhesive” pillars, no nuclear deformation was observed (0%). Strikingly, on “interspace adhesive” pillars, the nuclei were completely inserted between the pillars, resulting in a maximum of the nuclear deformation (100%). Altogether, these results show that adhesion to the lateral edges of pillars but not to the top of the pillars is necessary for driving nuclear deformation. Surprisingly, modification of pillar chemistry and topography did not affect hMSC nuclear deformation (**S5**), indicating that restricting adhesion to the lateral edges of the pillars is not sufficient to induce cell and nucleus deformation.

**Figure 4.**
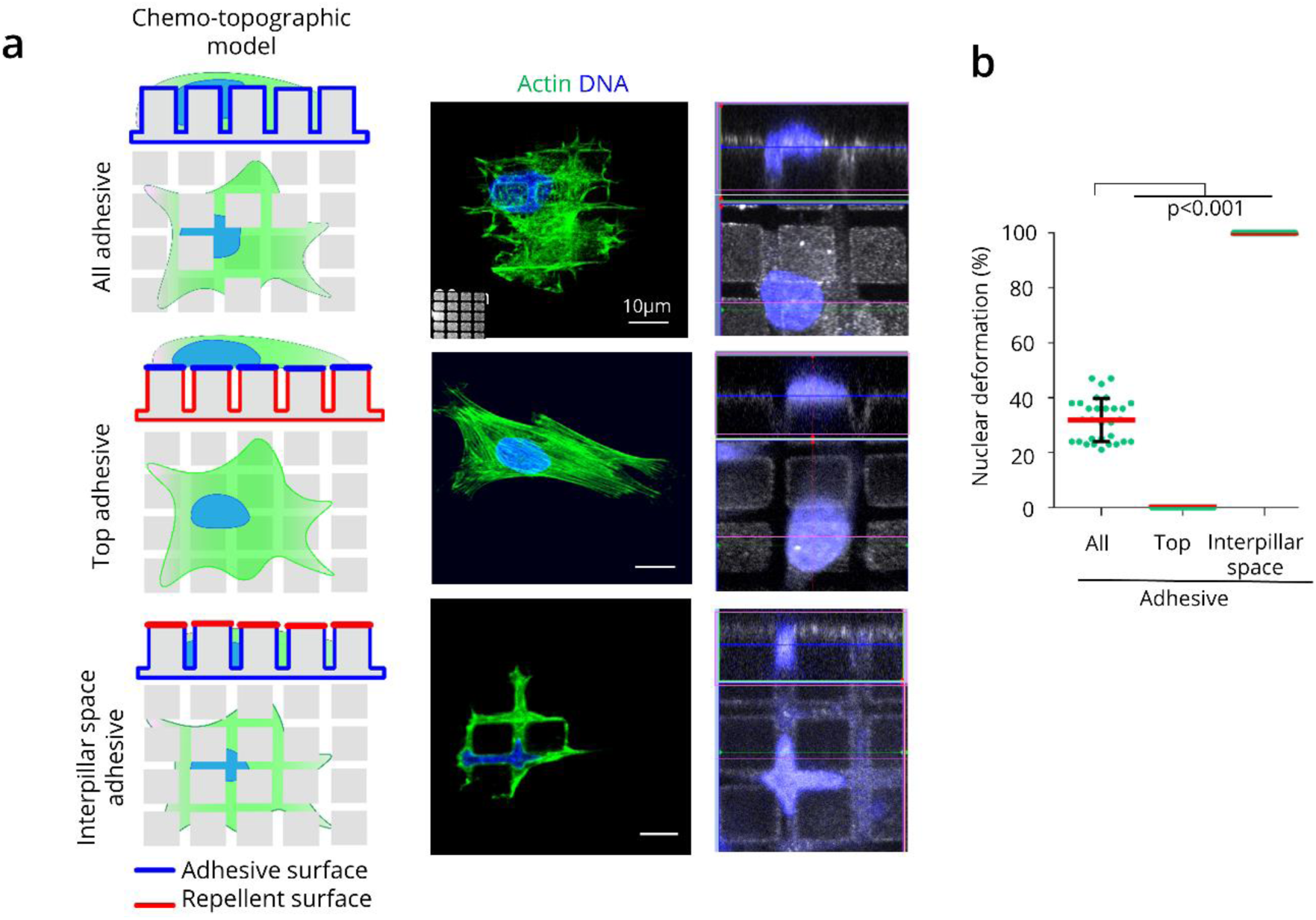
Effect of micropillars chemo-topographic modelling on nuclear deformation. **a)** Left, Sketch representing chemo-topographic alteration of 3/10 µm (space/height) micropillar surface chemistry (Blue: cell-adhesive PnBA coating, red: cell-repellent PDMA coating). Middle, Fixed cells stained for actin (green) and DNA (blue). Right, Orthogonal images of nuclei (blue) on 3/10 µm (space/height) micropillars. **b)** Quantification of nuclear deformation of SaOs-2 nuclei on different chemistry coated surfaces compared to ‘All adhesive’ surface; (n=30 nuclei/surface over three independent experiments).

### *In silico* analysis of nuclear stress during cell confinement

To attain further insight on the effect of micropillar topography on the mechanical stress on the nucleus during self-confinement, we used an *in silico* cell model published by Vassaux et al.(30) It assumes that the cell’s mechanical homeostasis results from the equilibrium between contraction forces generated by the actomyosin network and compression forces originating from microtubules, intermediate filaments and the cytosol. The LINC complex is modeled as connections between the nucleoskeleton and the three cytoskeletal networks (microfilament, microtubules and IF). Briefly, the nucleus was represented as a dense packing of particles in contact, enclosed by a stretchable envelope. The model represents the mechanical stress during nuclear deformation depending on variables such as presence and absence of: i) LINC ii) apical actin cap and iii) pillars dimensions and chemistry modulations. The mechanical stress is assumed to be caused by nuclear components themselves and the cytoskeletal elements (**Fig. 5, S6**). In the absence of the LINC complex, nuclear deformation and mechanical stress on the nucleus are reduced compared to the reference model comprising the LINC complex (∼4-fold and 2-fold less respectively) (**Fig. 5a-c**) which recapitulates the experimental data. This shows that in the absence of LINC complexes, the cytoskeleton is unable to transmit forces (observed by least mechanical stress) required for nuclear deformation. We modeled the pillar dimensions and chemistry constraints to quantify the forces on the nucleus. We used the “all adhesive” pillars as a reference for calculation. The mechanical stress experienced by the nucleus during deformation was at a minimum on “all adhesive” and “top adhesive” pillars, whereas it was at a maximum on “interspace adhesive” pillars, reaching twice the value observed with the reference surface (**Fig. 5d-f**). Taken together with the experiments, the highest mechanical stress observed on the interspace adhesive pillars configuration suggest that cells probably self-confine by pulling themselves on the substrate. Besides pulling forces, cells in confined spaces also experience pushing forces that can be generated by the apical actin cap.(31) Lastly, we examined the influence of actin cap on both nuclear deformation and mechanical stress. We observed that loss of apical actin has no impact on mechanical stress or deformation of nucleus which also coincides with the experimental results (**Fig. 5g-i, S7**), suggesting that the pushing down forces might not play a significant role in deforming the nucleus.

**Figure 5.**
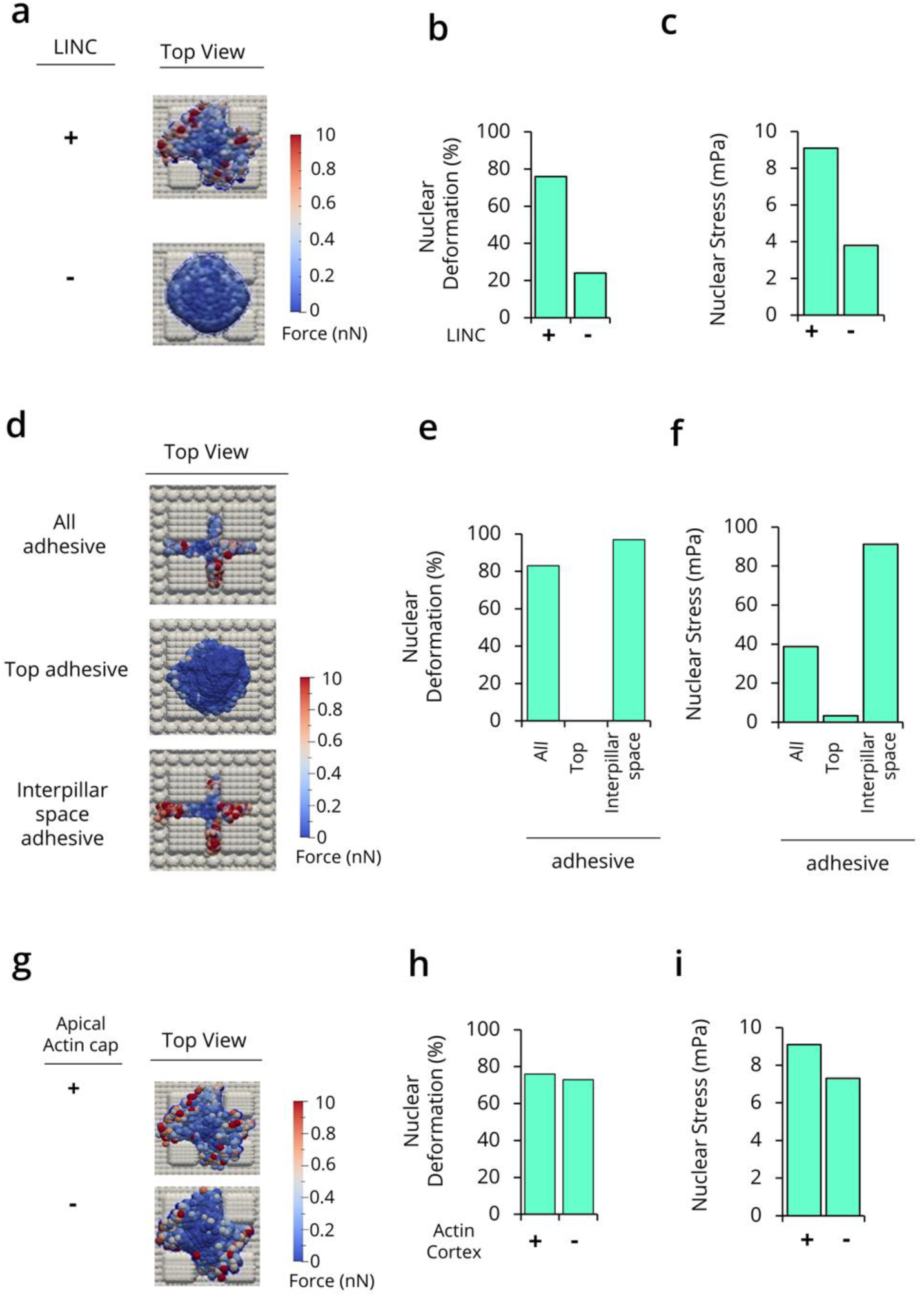
*In silico* cell model to analyze mechanical stress on nucleus during deformation. Computational model images showing the nucleus deformation and mechanical stress in mPa on pillars with and without LINC (**a-c**), after chemistry and topography surface modifications (**d-e**) and with and without LINC (**g-i**). Computational model images of adhesion on the pillars showing the nucleus deformation and overall stress. The values of the deformation of the nucleus obtained from the in-silico model simulations were calculated the same way as in vitro experiments using a cross-sectional view and dividing the area above pillars by the total area of the nucleus (see Fig. S1).

## DISCUSSION

The mechanisms by which cells exert forces to confine their cell body and particularly their rigid nucleus into narrow constrictions is not yet fully understood. We dissect here the role of different cytoskeletal elements during spontaneous nuclear confinement using a multidisciplinary approach that combines cellular and molecular biology techniques, 3D surface chemistry modifications and *in silico* cell modeling. We show that the nucleus is pulled down by the cell through its LINC complex using actomyosin contractility assisted by vimentin (Fig. 6).

**Figure 6.**
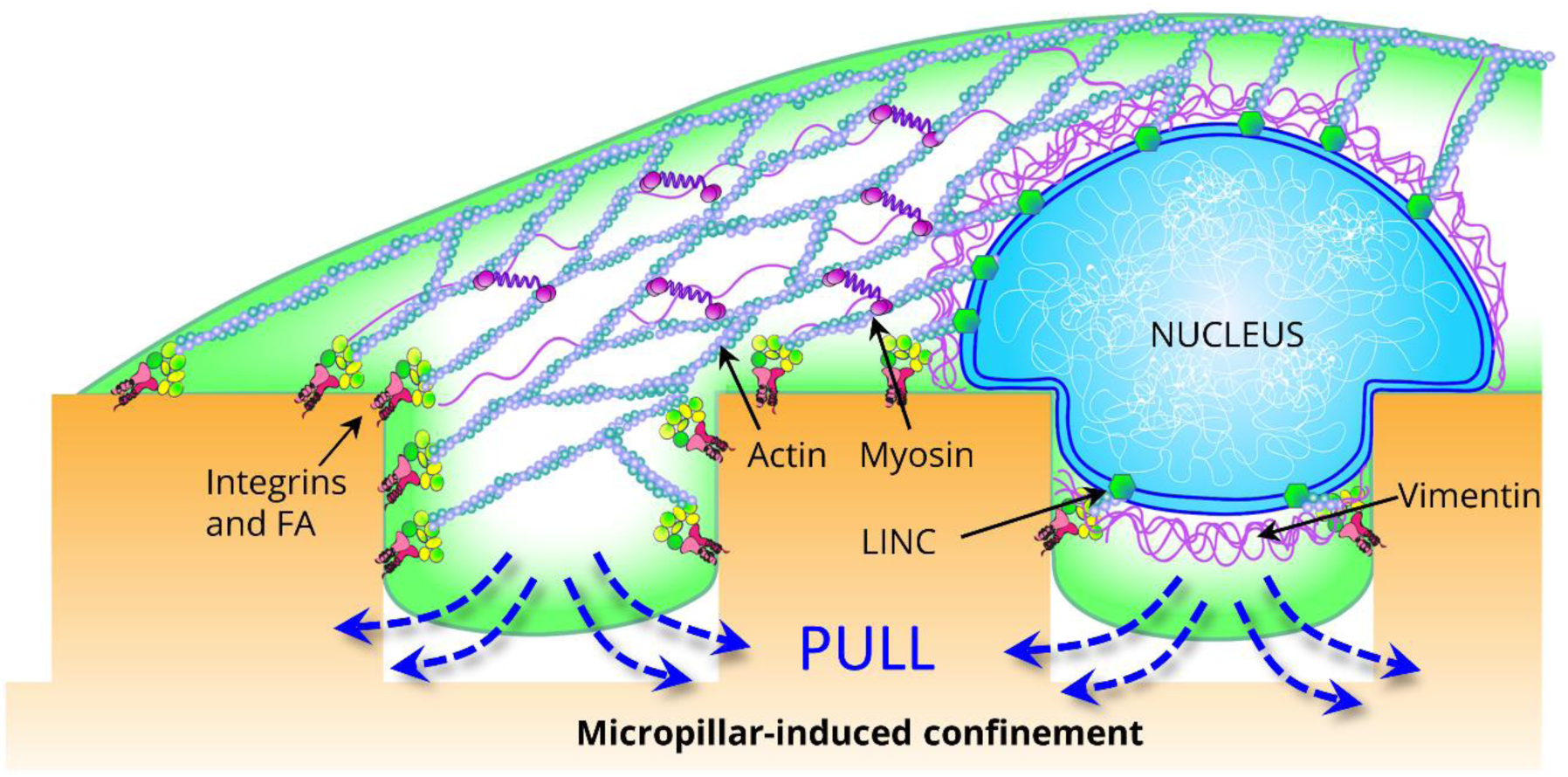
Proposed mechanism underlying nuclear deformation. Actomyosin pulling force, coupled to FAs and LINC complexes, drives cell-nuclear deformation on pillar topography. In this system, vimentin IF also assist in nuclear deformation.

### Micropillar arrays are a versatile tool to study nuclear deformation

Here we study nucleus deformation due to cell-generated forces using our micropillar-based self-confinement assay. This strong deformation of the nucleus is maintained for hours, yet it is reversible: as cells divide, they lose then regain their deformation. This assay allows a high degree of versatility and is easily tunable as we show in this manuscript. 1) We can easily change the size of the constrictions to modulate the extent of deformation. 2) We can change the surface chemistry to determine the directionality of forces. 3) We can rapidly add drugs and other chemicals to determine the actors responsible for the phenomenon. It is important to note that the phenomenon we study is not due to migration into the surface (migration can only occur perpendicularly to the surface). We are thus studying nuclear deformation by the cell specifically and decoupled from migratory cues. Nonetheless, these mechanisms may be used by the cell during cell migration to deform the nucleus.

### The forces on the nucleus involve the LINC complex, vimentin, actin and myosin contractility

We found that actin, myosin activity and vimentin are necessary for nucleus deformation and confinement by the cell. This is in agreement with studies that have linked migration through narrow constrictions with vimentin and actin activity (reviewed by McGregor et al.(3)). We demonstrate that disruption of the LINC complex using a dominant negative KASH construct results in severe loss of deformation of the nucleus. This strongly indicates that the LINC complex is involved in exerting force on the nucleus during the deformation. These results reflect studies in which the LINC complex is involved in moving the nucleus in 2-D (26), nesprin-2 knock-down increases transit time through narrow constrictions (32) and a linker of actin and the LINC complex is necessary for nucleus translocation during migration through narrow constrictions.(33) The nesprins that are likely to exert forces on the nucleus are the two giant isoforms that bear actin-binding domains, however we found that knocking down these isoforms individually did not affect nucleus deformation. This may be due to redundancy between the two giant nesprins, or may indicate that other nesprins could contribute. Indeed, Nesprin-3α can aid force transmission to the nucleus through vimentin.(35) Further study may thus be necessary on the individual nesprins and intermediary proteins involved in this deformation.

### Cells pull on their nucleus

Cells that only adhere to the tops of pillars are unable to deform their nuclei and the extent of deformation is strongly increased when adhesion is constrained to the sides of the pillars (fig 4). Thus, contractile forces from above the pillars are not required to deform the nucleus. This striking result does not agree with proposed mechanisms by which nucleus deformation during migration through narrow constrictions is caused by contraction at the rear of the cell.(3) Other studies have suggested that the nucleus is being pulled during migration (3), also based on the shape of the nucleus as it is deformed.(15) It is important to note that here we study the deformation of the nucleus by the cell decoupled from migratory cues, and thus contraction at the rear of the nucleus may be necessary for forward migration of the cell but not for nuclear deformation. It is therefore likely that the mechanisms are dependent on the particular circumstances of the nuclear deformation, but multiple mechanisms may also be involved, which may synergistically contribute to forward migration through constrictions.

### The forces exerted on the nucleus are generated on the lateral sides of the pillars

Intriguingly, forces exerted from the sides of the pillars are sufficient to pull on the nucleus: there are no focal adhesions at the bottom of the pillars and the cells deform extensively in tall pillars (10 µm, fig. 4). Our observations of the cytoskeletal architecture and the progression of focal adhesion formation indicate that, as focal adhesions form on the edges of the pillars, the nucleus is pulled down by the actin cytoskeleton, aided by intermediate filaments. The progression of the nucleus between the pillars thus closely follows the appearance of focal adhesions, suggesting closely-related mechanisms for cell adhesion and nucleus pull-down, and may indicate direct coupling between focal adhesions and the LINC complex.

### Lamin A/C does not affect the extent of deformation

Previous studies have found that cells with reduced lamin A/C levels translocate their nucleus through narrow constrictions more quickly than their wild-type counterparts. (36, 37) We found that increasing or decreasing nuclear deformability by reducing or increasing lamin A/C expression does not affect the extent of deformation by the cell at 24 hours. However, future studies should focus on whether lamin A/C levels may affect the speed of deformation on micropillars, which would be in agreement with results obtained in the migration studies.

### The mechanisms by which healthy cells prevent deformation

We had previously shown that MSCs react to the pillars by spanning the top of the pillars rather than adhering between the pillars and deforming the cell body and nucleus. We show here that even in circumstances in which the adhesion to the top of the pillars is prevented, the MSCs adhere in between the pillars without deformation. Thus MSCs likely have mechanisms to prevent the deformation of the nucleus. We observed that the A-type lamin levels were comparable in between the SaOs-2 cells and MSCs, indicating that lamins are not the sole determinants of nuclear deformation. The difference may thus lie in the coupling between the nucleus and the cytoskeleton, or the manner in which the cells engage contractility at the LINC complex. Direct mechanical coupling between adhesions and the nucleus may be lacking in these cells.

## CONCLUSIONS AND OUTLOOK

Our 3D micropillar array platform provides a simple highly tunable approach to access nuclear deformation using cell-generated forces decoupled from migration. We have already shown nuclear deformation in a variety of cell types.(20-23) Recently the nuclear deformability of human pluripotent stem cells during early germ layer specification was investigated using 3D micropillar array platforms and found to be lost during ectoderm differentiation.(38) Nuclear deformability on micropillars could thus be used to determine differentiation. This technique could be further exploited to explore the 3D organization of chromosome territories and gene expression pattern modifications and unravel other molecular players/cell signaling pathways related with confinement-induced nuclear deformation.

## DATA AVAILABILITY

The processed data required to reproduce these findings are available to download from http://dx.doi.org/10.17632/psfmnh8n2y.1. The raw data are available upon request.

## CREDIT AUTHOR STATEMENT

N.T.W. Conceptualization, Data curation, Formal analysis, Investigation, Methodology, Writing - original draft; S.A. Investigation, Methodology; F.B. Investigation, Methodology; M.E. Investigation, Methodology; I.B. Investigation, Methodology, Validation; T.P. Investigation, Methodology, Validation; M.V. Investigation, Methodology, Software; J-L.M. Methodology, Supervision, Validation; J-N.F. Writing - review & editing; J.R. Funding acquisition, Methodology, Supervision; P.M.D. Conceptualization, Methodology, Writing - review & editing; L.P. Conceptualization, Methodology, Supervision, Writing - review & editing; K.A. Conceptualization, Funding acquisition, Supervision, Writing - review & editing.

## ACKNOWLEDGEMENTS

We are grateful to Dr. Nicholas Borghi (Jacques Monod Institute, Paris-France) for providing DN-KASH mcherry plasmid and Dr Tanmay Lele (University of Florida, USA) for Lamin-A GFP plasmid. This work was supported by Region Alsace [NTW and FB PhD grants], Deutsche Forschungsgemeinschaft [SA and ME PhD grants], IRTG Soft Matter [NTW, SA, ME, PMD], La Ligue contre le Cancer [REMX17751 to PMD and CCIR-GE120193 to KA] and the Fondation ARC [PDF20161205227 to PMD].

## SUPPLEMENTARY FIGURES

**S1:**
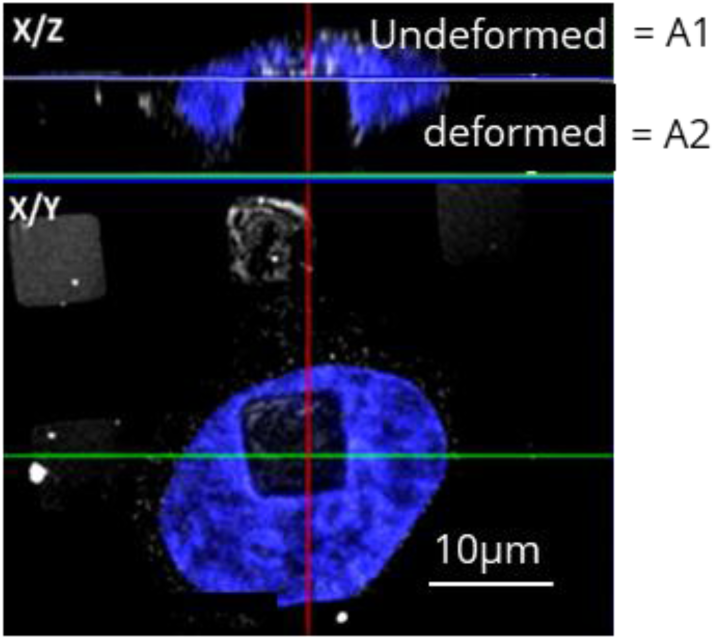
Calculation of nuclear deformation. Image shows the cross sectional area of nucleus (blue) on micropillar surface. The area above (A1) and below (A2) pillar is calculated using ImageJ/Fiji. The deformation of nucleus is calculated in percentage by the formula A2/A1+A2*100.

**S2:**
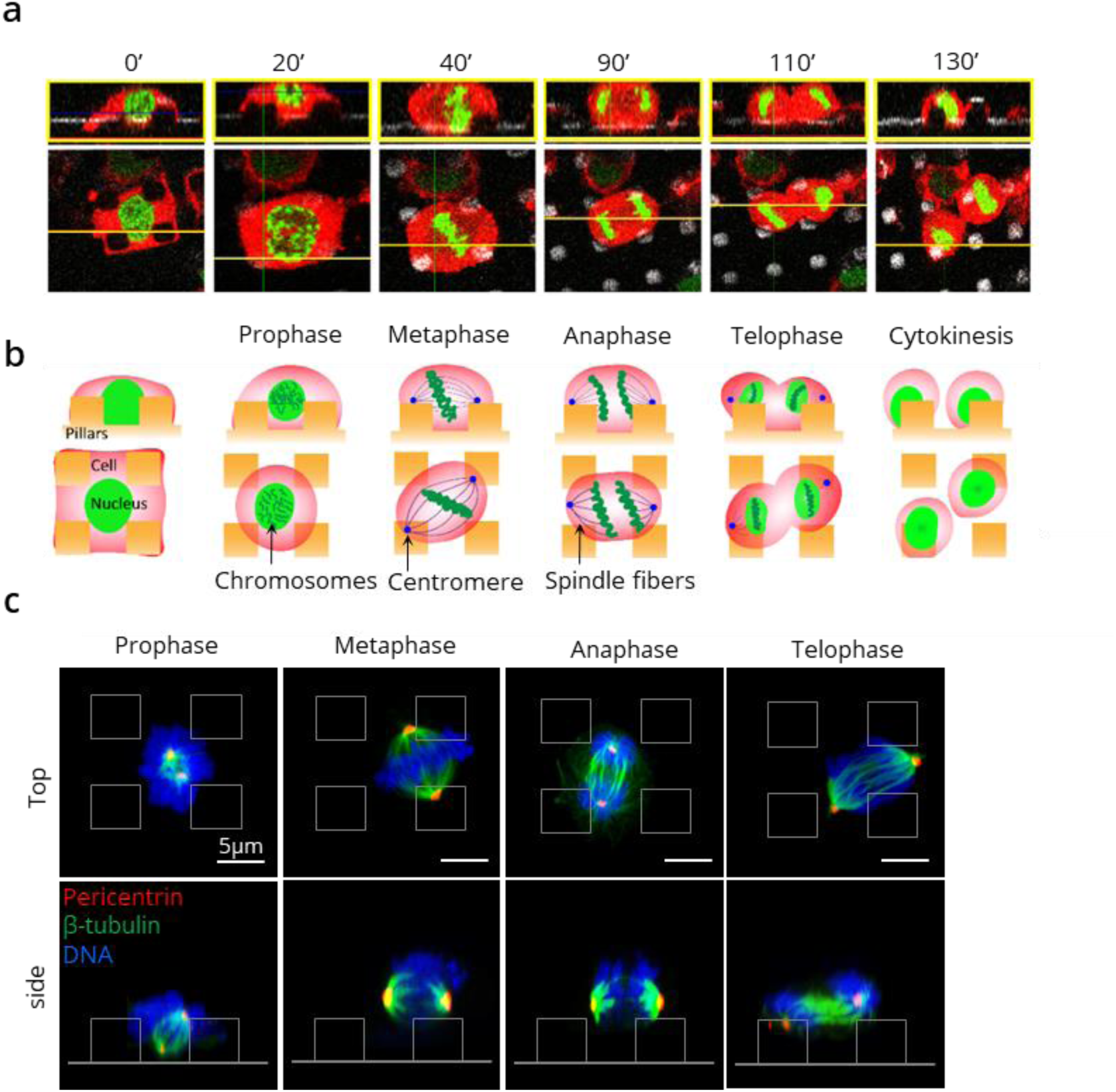
Mitosis on pillars. **a)** Orthogonal view of the SaOS-2 cell dividing on the micro-pillars with time, transfected with histone actin-RFP and histone-GFP. **b)** The drawing illustrates the mitotic phases performed by SaOS-2 on pillars. **c)** Confocal z stack projection images showing mitotic phases and centromere position during division (pericentrin-red, β-tubulin-green and DNA-blue).

**S3:**
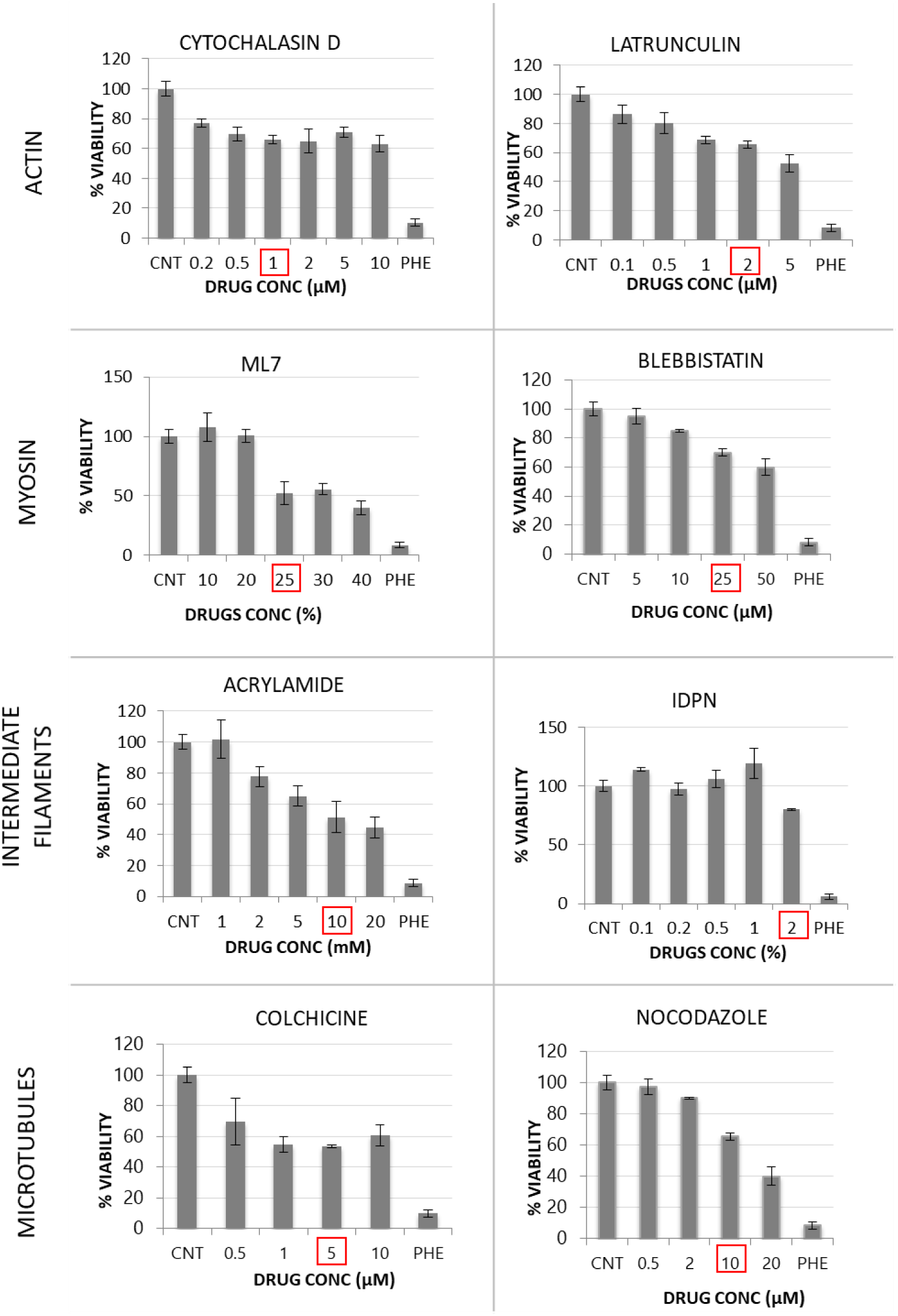
MTT assay. Graphs showing the MTT viability test and the respective concentrations chosen (red box) from different concentrations of drugs after 6 hours. Error bars represent ±SD resulting from three replicates over three independent experiments.

**S4:**
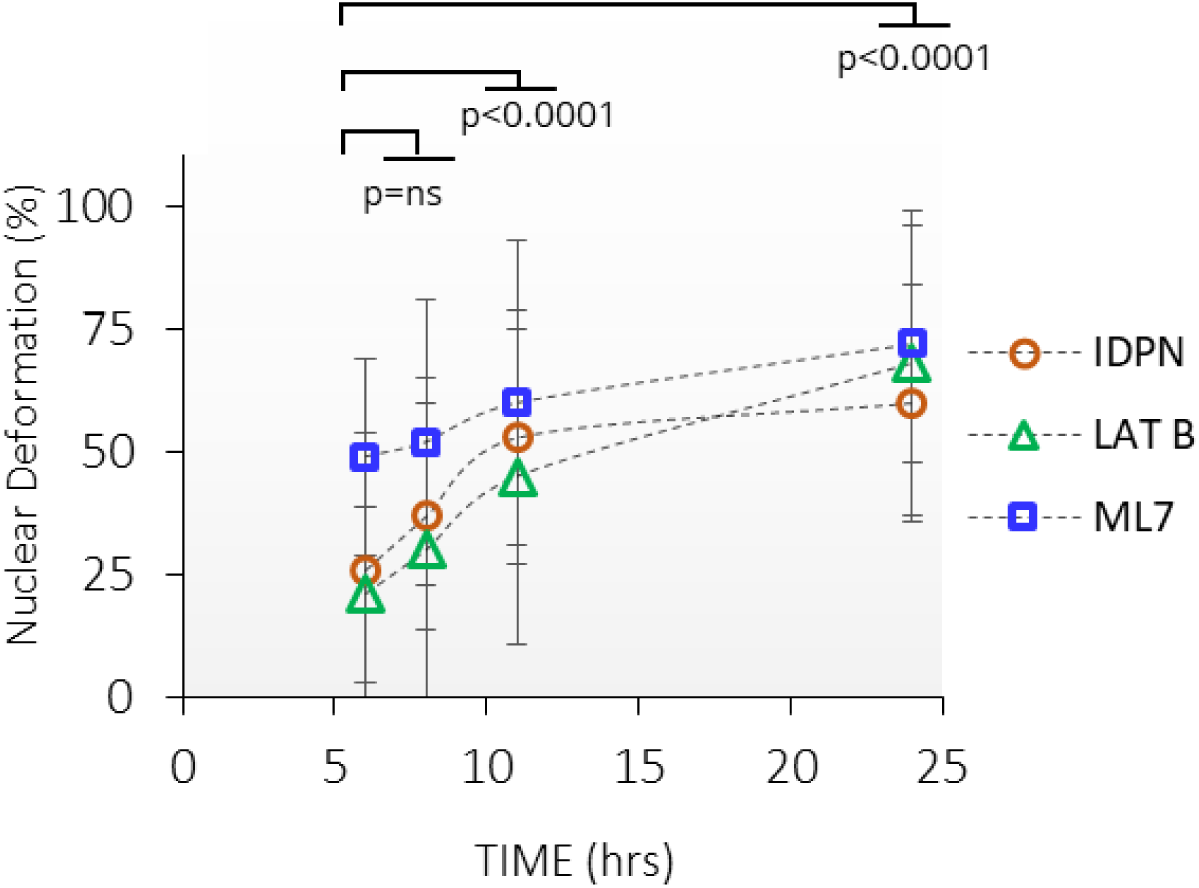
Nuclear deformation is restored after drug washout. Graph showing the retention of nucleus deformation at 2, 5 and 20 hours after drug washout. The symbols show mean nuclear deformation percentage at each time point (n=30/ drug treatment), where ‘n’ is number of cells analyzed from three independent experiments. The symbols represents the drugs (circle=IDPN; triangle=Lat B and square= ML7). Error bars represent ±SD. Red dotted line represents drug washout after 6 hours of initial drug treatment. P-values were calculated by unpaired two-tailed student’s *t* test comparisons test for comparison.

**S5:**
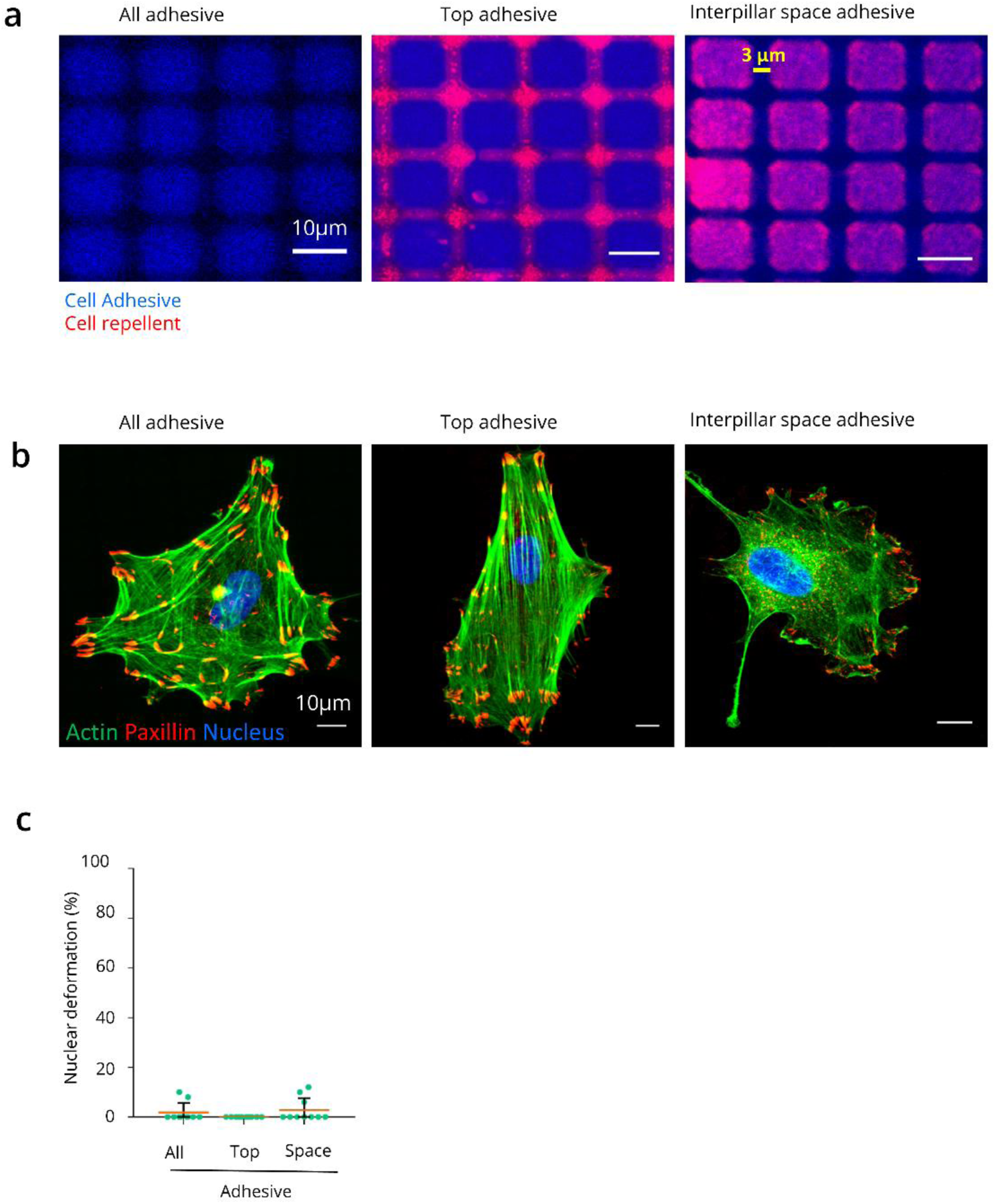
Validation of chemistry coatings and hMSC behaviour on chemo-topographic variations. **a)** Images showing modified surfaces: blue-cell adhesive and red-cell repellent surface. **b)** hMSC cells on modified surfaces, immunostained for actin-green, paxillin-red and nucleus-blue. **c)** Graph showing nuclear deformation percentage of hMSC nuclei on different chemistry coated surfaces; (n=30 nuclei/surface), where ‘n’ is number of cells analyzed from three independent experiments with mean (red line). All error bars represent ±SD.

**S6:**
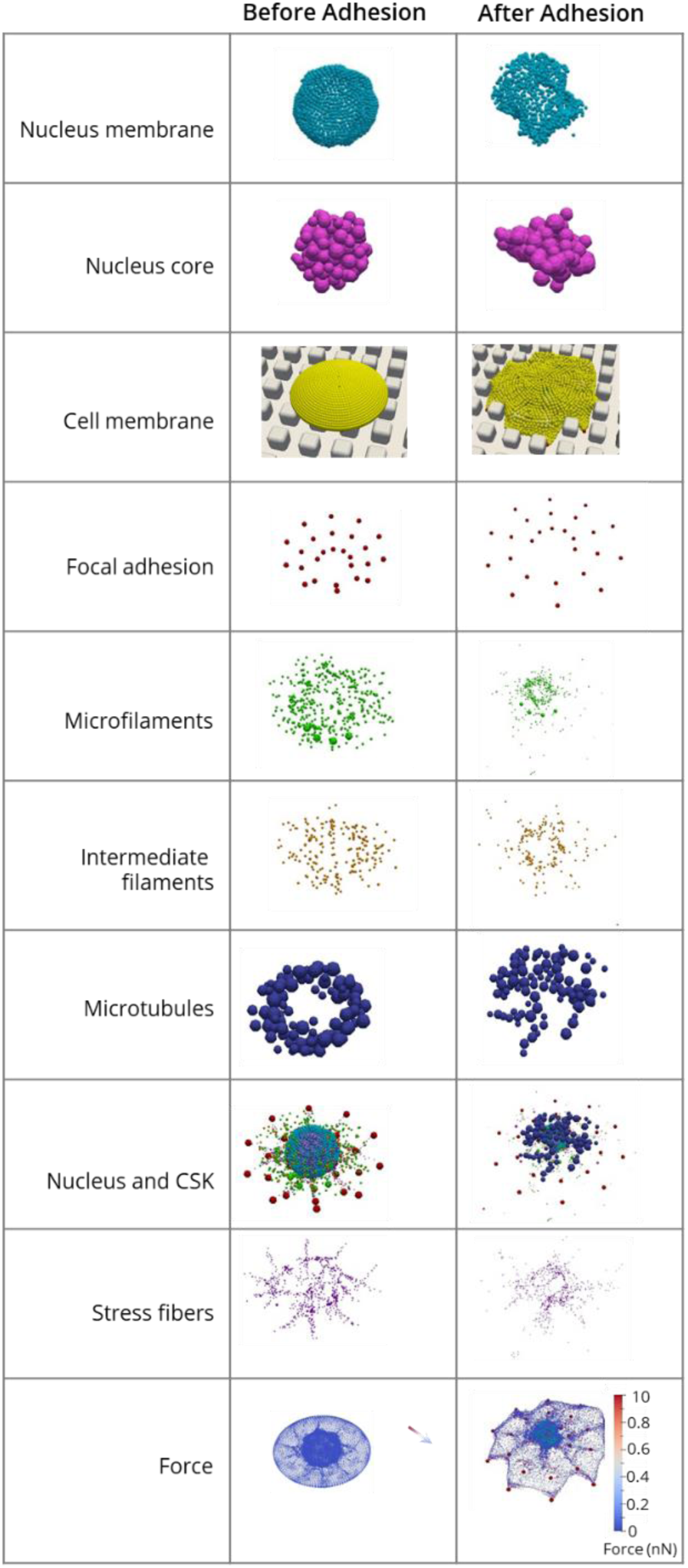
Figure showing elements used in in-silico cell model simulation. Different elements used in in silico model simulation to calculate mechanical stress on nucleus during deformation.

**S7:**
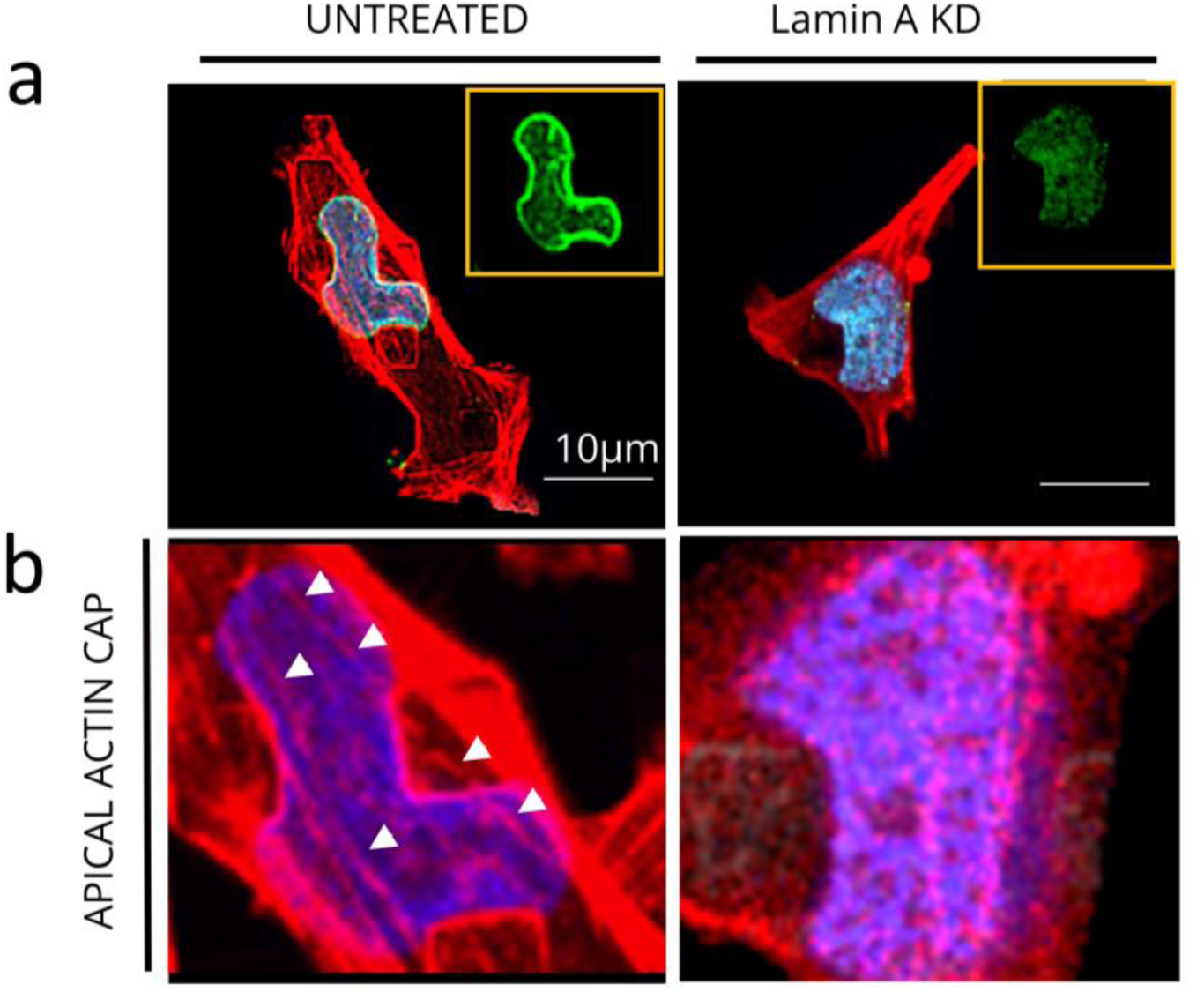
Actin cap reduction by lamin A KD. **a)** SaOs-2 cells on pillars of lamin A (green) and actin (red) in knockdown and untreated control cells after 24 hrs. **b)** Zoom images showing apical actin cap fibers in untreated cells and disrupted apical actin cap fibers after lamin A KD.

**S8:**
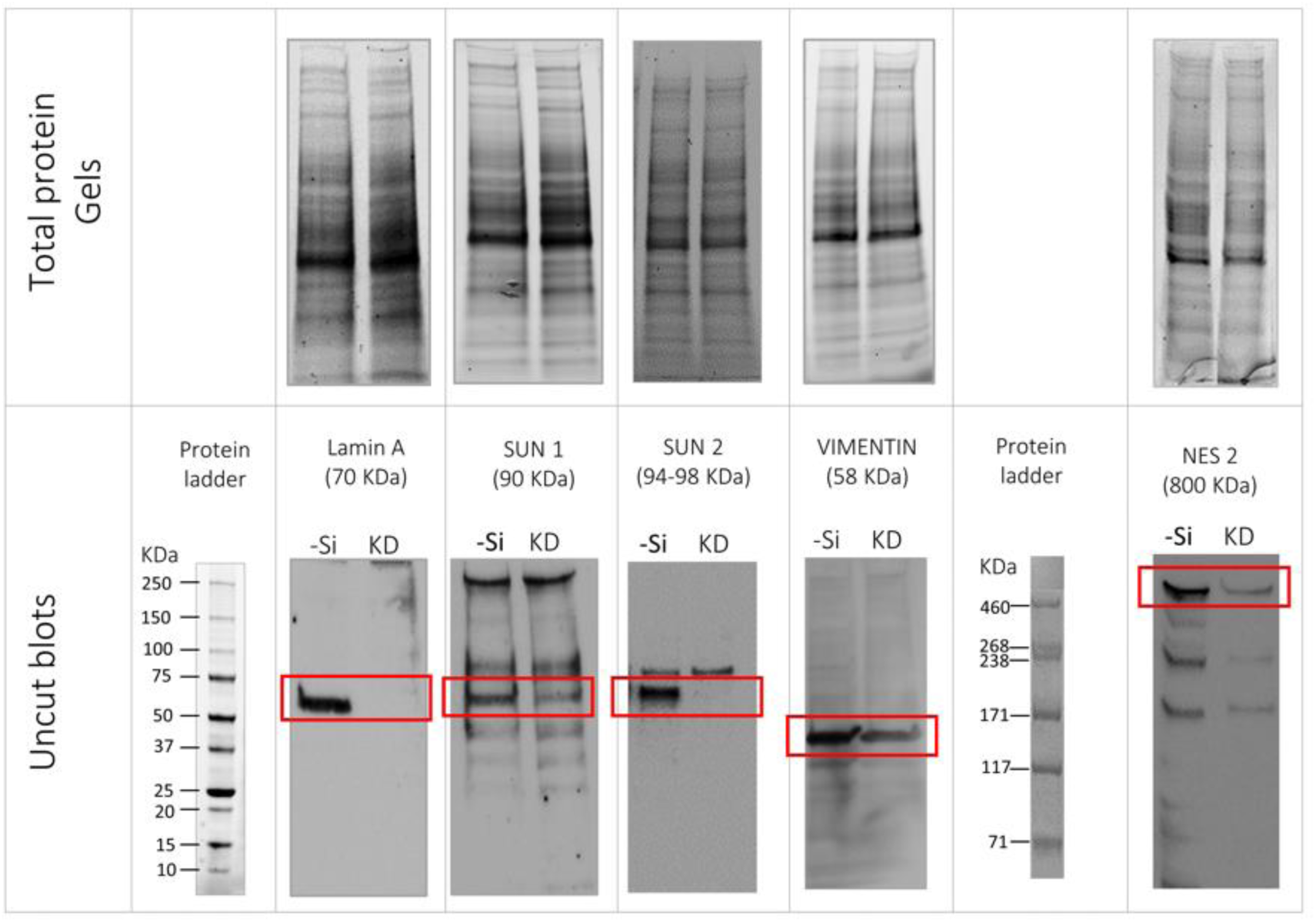
Western blot. Total protein gel images for KD of lamin A, SUN1, SUN2, nesprin 2 and vimentin used for protein normalization and corresponding immunoblots.

**S9:**
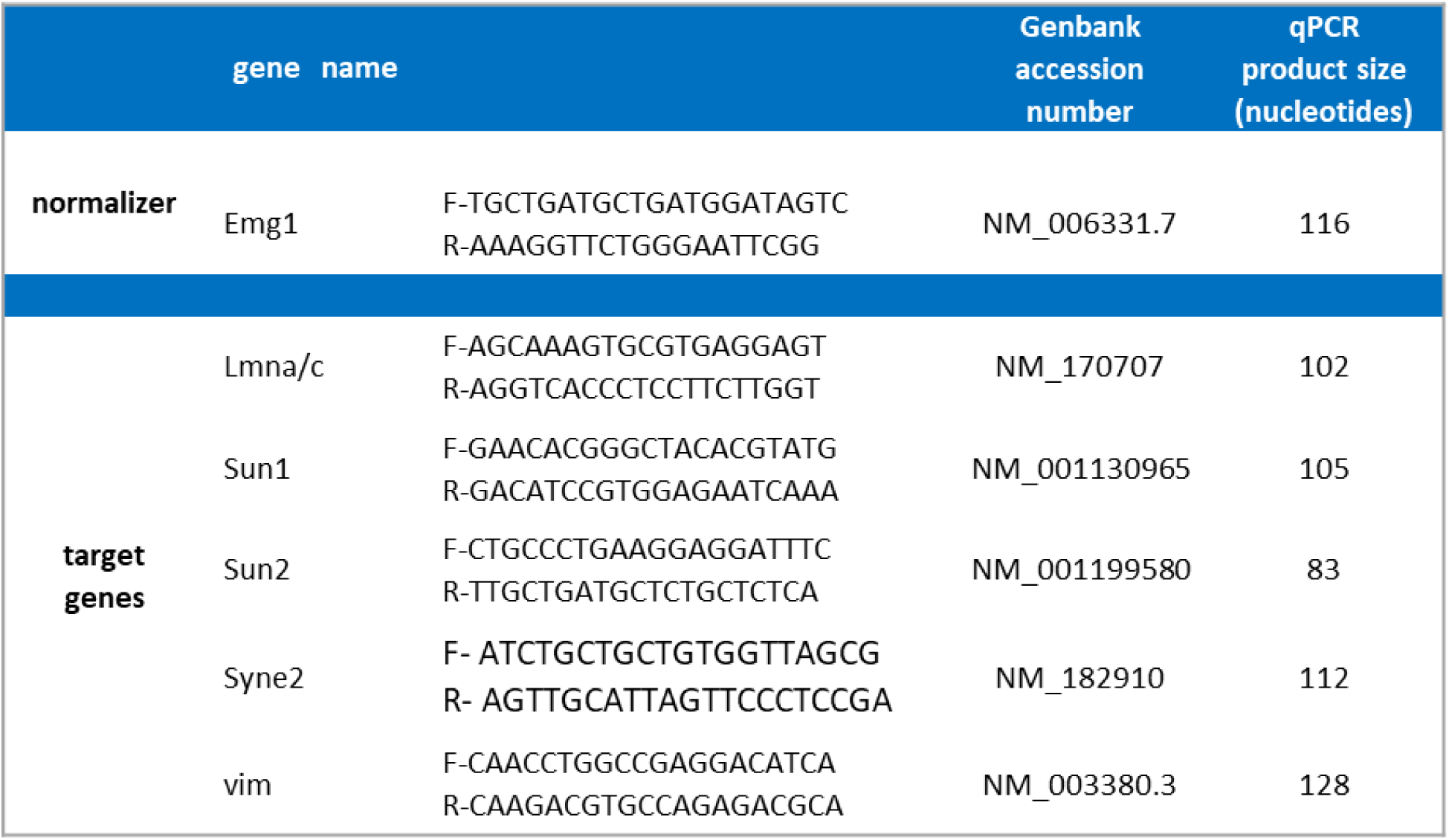
qRTPCR primer and cycle information. Nucleotide sequences of pairs of primers used for qPCR experiments. “F” and “R” indicate forward and reverse primers respectively. Emg1 (18S methyltransferase), Lmna (lamin A/C), Lmnb1 (lamin B1), Sun1 (Sad1 and UNC84 domain containing 1), Sun2 (Sad1 and UNC84 domain containing 2), Syne2 (nesprin 2), vim (vimentin).

## Legends for the video files

**SM1: Adhesion and deformation**

Movie showing the deformation of a cell (red) and its nucleus (green) over time. Nuclear deformation increased with time. At around 1 hour the deformation is initiated and seems to be completed at about 6-8 hours. At 12 hours the cell spreads more on the nearby pillars.

**SM2: Mitosis on pillars**

Cell dividing on pillars with time. The dividing cell ascends up the pillar structure to perform cell division. Mitosis is performed above the pillar structure until the daughter cells are formed, which immediately descend back in between the pillars and deforms immediately after division.

